# An ancestral role of SIAMESE-RELATED proteins in cell-division arrest predates the emergence of endoreduplication in land plants

**DOI:** 10.64898/2026.11.23.768773

**Authors:** Facundo Romani, Marius Rebmann, Ignacy Bonter, Jim Haseloff

## Abstract

- Endoreduplication is central to cell enlargement and differentiation in flowering plants, yet its evolutionary origin and relationship to core cell-cycle regulators remain poorly understood. SIAMESE-RELATED (SMR) proteins promote endoreduplication in angiosperms, but their ancestral function outside seed plants is unknown. We investigated the evolution of endoreduplication across land plants and the physiological role of SMRs in the liverwort *Marchantia polymorpha*.
- Ancestral-state reconstruction of endoreduplication levels in the vegetative body of the dominant generation supports the absence of this trait in the common ancestor of land plants, with at least two independent evolutionary origins in mosses and seed plants.
- Phylogenetic analysis of SMR proteins supports their origin before the diversification of land plants and their presence in species that lack endoreduplication.
- In Marchantia, SMR acts as a CDK inhibitor by suppressing cell division and promoting cell expansion, while loss of SMR results in increased growth but does not impair cell differentiation. In addition, an SMR gene from the alga *Chara braunii* is functional in Marchantia, revealing deep conservation of SMR activity.
- These findings are consistent with SMRs having originated before the emergence of land plants as regulators of cell size and suggest that their role in promoting endoreduplication is a derived function.

## INTRODUCTION

Endoreduplication is a modified cell cycle which involves multiple S-phases without mitosis, resulting in increased ploidy. This cell-cycle variant occurs across diverse eukaryotic lineages and may have evolved repeatedly across eukaryotes (Lee et al., 2009). In flowering plants, endoreduplication plays an important role in the growth of epidermal tissues (Melaragno et al., 1993) and in the differentiation of trichomes (Churchman et al., 2006), trichoblasts (Li 李磊 et al., 2025) and giant cells in sepals (Roeder et al., 2010; Lang and Schnittger, 2020). Endoreduplication results in endopolyploidy, which can be detected by flow cytometry as discrete peaks corresponding to integer multiples of the haploid or diploid genome content. Endoreduplication is common in flowering plants and mosses (Bainard et al., 2020), involving a large proportion of vegetative cells with high endoploidy levels. Outside of flowering plants, endoreduplication is absent or rare in several lineages (Bainard et al., 2011; Bainard et al., 2013; Bainard and Villarreal, 2013; Nishihama et al., 2015). However, a consistent phylogenetic mapping of this trait is lacking, leaving its evolutionary origin unresolved.

In plants, endoreduplication is associated with cyclin-dependent kinase inhibitors (CKIs). Beyond endoreduplication, CKIs constitute a central layer of eukaryotic cell-cycle control by modulating the activity of cyclin-CDK complexes and integrating developmental and environmental signals by promoting cell-cycle arrest (Peres et al., 2007; Yi et al., 2014; Cao et al., 2018; Kimata et al., 2020). In mammals, members of the Cip/Kip family (p21, p27, p57) inhibit CDK activity and thereby coordinate proliferation with differentiation and stress cues (Kimata et al., 2020). This function is conserved in plants, where the homologous KIP-RELATED PROTEINS (KRPs) inhibit Cyclin D-CDKA complexes (Wang et al., 1997; De Veylder et al., 2001). In addition to KRPs, plants possess the SIAMESE-RELATED proteins (SMRs), a plant-specific CKI family that emerged independently of Cip/Kip proteins.

In flowering plants, SMRs are best known for promoting the entry into the endocycle by inhibiting CDKs, thereby coupling cell-cycle arrest to cell enlargement and differentiation (Churchman et al., 2006). SMRs were first identified through the *siamese* (*sim*) mutant, which revealed a gene controlling endoreduplication and inhibiting mitotic cell division during trichome formation (Walker et al., 2000). Plants defective in SIM display ectopic cell divisions in the unicellular trichomes of *A. thaliana,* whereas SIM overexpression causes cell enlargement (Churchman et al., 2006). However, because functional studies of SMRs have been largely restricted to flowering plants, it remains unclear whether their association with endoreduplication reflects an ancestral property of the family or a derived function acquired during plant evolution.

Studying the molecular mechanisms associated with CKIs in diverse plant species could help to answer fundamental questions about the evolution of the cell cycle. The liverwort *Marchantia polymorpha* (hereafter Marchantia) is a powerful model for evolutionary cell biology because it features a reduced complement of cell-cycle regulators (Romani et al., 2026) and lacks vegetative endoreduplication (Nishihama et al., 2015). The single KRP gene in Marchantia, MpKRP, regulates cell size through inhibition of cell division (Romani et al., 2026), supporting the idea that CKIs can regulate growth independently of endoreduplication. In the model moss *Physcomitrium patens*, PpSMR12 inhibits CDKA in vitro and complements the Arabidopsis *siamese* mutant (Kumar et al., 2015), indicating that SMR biochemical activity is conserved outside flowering plants. However, the endogenous physiological function of SMRs in bryophytes remains largely unexplored (Hernandez-Garcia et al., 2021).

Here, we reconstructed the evolutionary history of endoreduplication across land plants and investigated the origin of the SMR protein family. We found that endoreduplication in the vegetative body may have emerged independently in mosses and seed plants, whereas SMR genes were already present in streptophyte algae. We found that the single-copy SMR gene in Marchantia, MpSMR, inhibits cell division and promotes cell expansion, similar to the function of MpKRP (Romani et al., 2026) and of SMR orthologues in angiosperms (Yamada et al., 2022). Loss-of-function mutants reveal that MpSMR modulates growth but is not required for cell differentiation. Furthermore, heterologous expression of an SMR from the charophyte alga *Chara braunii* in Marchantia phenocopies the cell enlargement phenotype observed in MpSMR. Overall, SMR genes predate the emergence of endoreduplication in mosses and angiosperms, supporting an ancestral function in cell-size control.

## RESULTS

### Endoreduplication is absent from the vegetative body of early-diverging land plant lineages and has independent origins in mosses and seed plants

Because endoreduplication varies among tissues and generations, we scored a single, comparable character across lineages: endoreduplication in the vegetative body of the dominant generation, the gametophyte in bryophytes and the sporophyte in vascular plants. Thus, to reconstruct the evolutionary history of endoreduplication across land plants, we compiled endoreduplication indices (EI) from published flow-cytometry surveys for 202 taxa spanning liverworts, hornworts, mosses, lycophytes, ferns, gymnosperms and angiosperms.

Values of EI below 0.1 were treated as absence of endoreduplication, because values in this range may reflect G2 nuclei rather than true endoreduplication (Bainard and Newmaster, 2010). For liverworts, hornworts, lycophytes, ferns, and gymnosperms, the endoreduplication indices indicate an absence of endoreduplication in every surveyed species (Figure 1). Among mosses, endoreduplication was observed in most major orders. Previous studies showed that Sphagnum species lack endoreduplication (Bainard and Newmaster, 2010). Interestingly, the early-diverging moss *Takakia lepidozioides* (Hu et al., 2023) also lacked endoreduplication. In angiosperms, endoreduplication was observed across several major groups sampled, spanning most major eudicot orders (including Brassicales, Fabales, Fagales, Caryophyllales, Rosales, Ranunculales, and Solanales) and monocots (Figure 1). Among the sampled representatives of Asterales and Lamiales, none showed EI values above 0.1. These observations indicate that endoreduplication is a widespread, though variable, feature of the angiosperm sporophyte.

**Figure 1.**
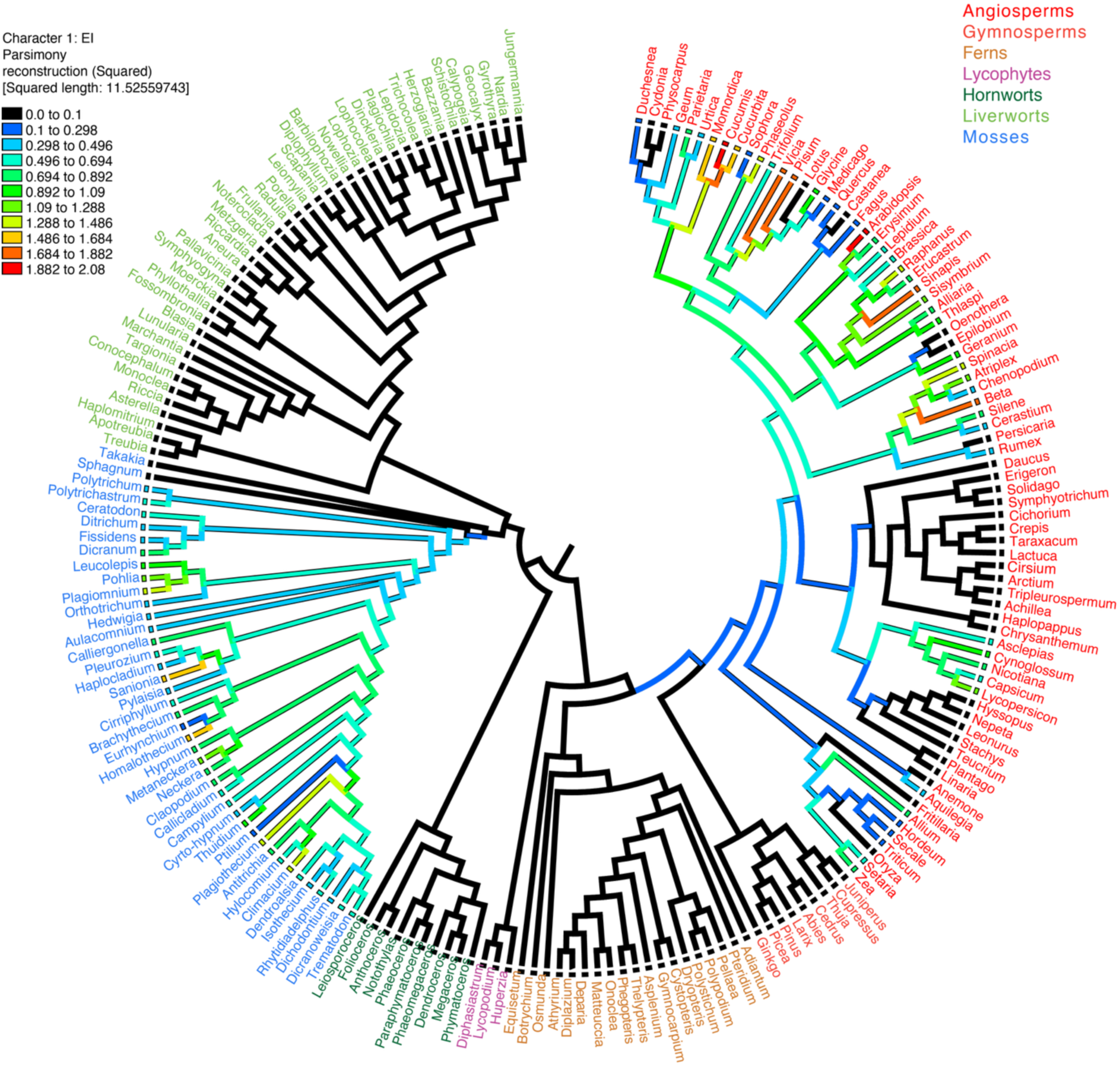
Evolution of endoreduplication across land plants. Phylogenetic distribution of endoreduplication index (EI) across 202 land-plant taxa, based on published flow-cytometry surveys. EI was scored for the vegetative body of the dominant generation: the gametophyte in bryophytes and the sporophyte in vascular plants. Branch colours indicate ancestral EI values reconstructed using squared-change parsimony (colour scale at left), while taxon labels are coloured according to major land-plant lineages (key at right). The complete tree is available at Supplemental File 1.

Squared-change parsimony reconstructed the ancestral EI of embryophytes below the threshold for endoreduplication (Figure 1). The same result was recovered for the ancestors of bryophytes and tracheophytes, both reconstructed below the threshold for endoreduplication. These results support at least two independent gains of endoreduplication across the land plant tree, one within mosses, after the divergence of Sphagnum (Bainard et al., 2020), and another along the seed-plant lineage. The reconstructed EI at the seed plant ancestor was only marginally above the threshold (0.1009), whereas the angiosperm ancestor showed a substantially higher value (0.1967).

To assess the robustness of this inference, we also reconstructed ancestral states treating endoreduplication as a binary character (EI ≥ 0.1). Maximum-likelihood reconstruction inferred three independent gains within angiosperms, marginal-probability reconstruction supported a single origin within the crown of seed plants. Despite these differences, all reconstructions agreed that endoreduplication was absent from the ancestors of embryophytes, bryophytes and tracheophytes, and that its evolution within mosses and seed plants occurred independently.

### SIAMESE-RELATED proteins predate the evolution of land plants

To investigate the evolutionary origin of SIAMESE-RELATED (SMR) proteins, we searched for SMR homologues across a broad sampling of plant genomes representing chlorophyte and streptophyte algae, bryophytes, lycophytes, ferns, gymnosperms and angiosperms. SMRs are short, intrinsically disordered proteins with limited primary-sequence conservation, which makes deep homology detection challenging. We therefore used the InterPro hidden Markov model for the SMR domain (IPR040389) as a conservative approach to identify candidate sequences and retained only high-confidence matches (E-value < 1e-5) for assessing presence/absence analyses and phylogenetic reconstruction. Only high-confidence matches (E-value < 1 × 10⁻⁵) were retained to maximize the specificity of the hits. All putative SMRs for Selaginella, Physcomitrella and Arabidopsis were retrieved, except for the highly divergent Arabidopsis SMR11/16 (Kumar et al., 2015).

SMR homologues were detected in all sampled land-plant lineages, from liverworts and hornworts to mosses, ferns, and angiosperms (Figure 2A). Among the streptophyte algae sampled, we recovered one clear SMR homologue each in *C. braunii* (Cb*SMR*) and *Klebsormidium nitens*. We also surveyed other available transcriptomes (One Thousand Plant Transcriptomes, 2019) and found an additional putative SMR homologue in *Nitella mirabilis*. Together, these findings indicate that SMR genes were already present in streptophyte algae before the colonisation of land. No clear homologues were detected in Zygnematophyceae. Given the short length and rapid divergence of SMRs, we interpret these data as a limitation of homology detection or secondary loss, rather than evidence of true absence.

**Figure 2.**
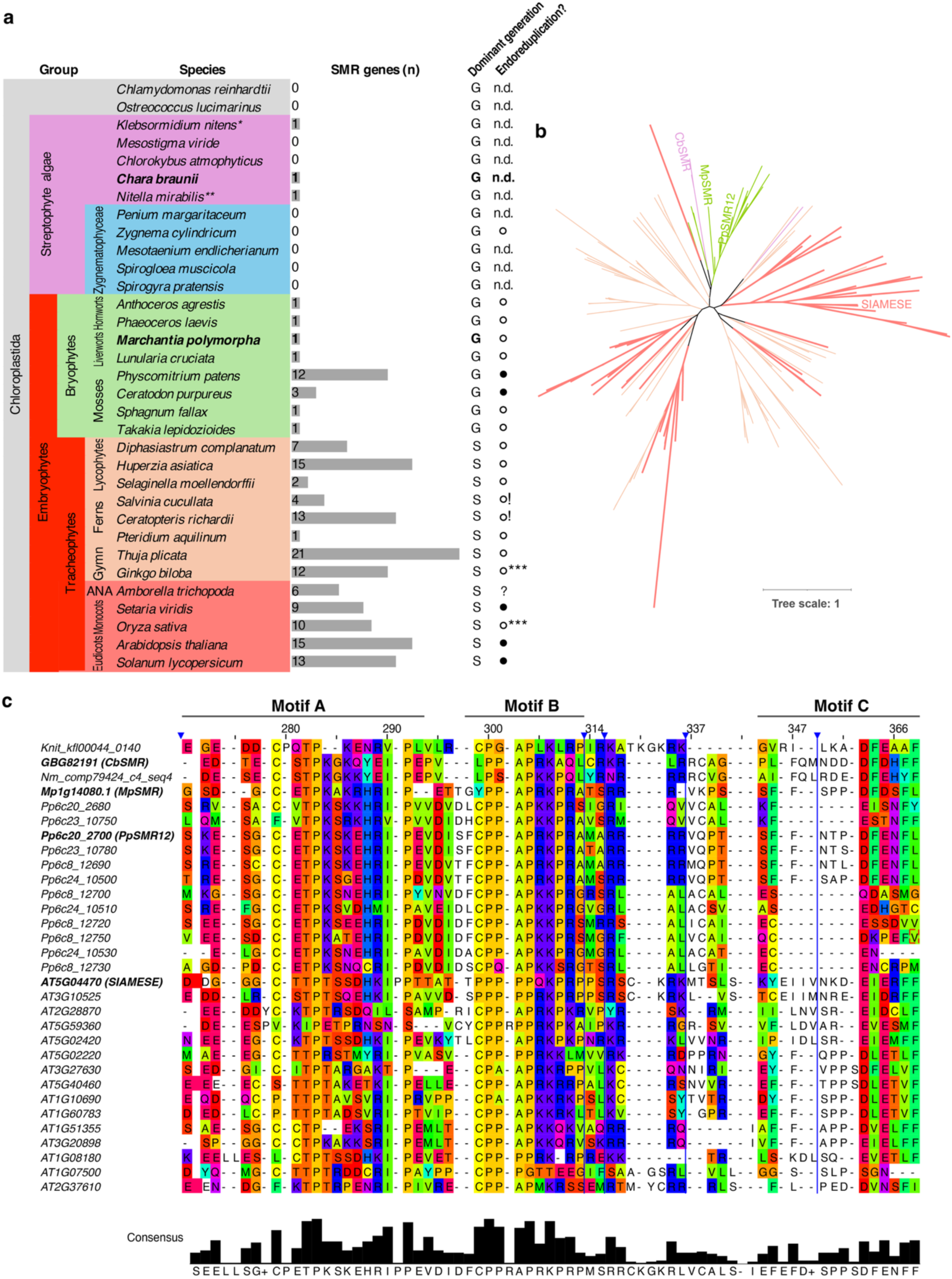
Evolutionary history of the SIAMESE-RELATED (SMR) protein family. (a) Summary of SMR homologues identified across streptophyte algae and land plant genomes, including their taxonomic distribution, presence or absence of endoreduplication, and copy number. * Indicates species represented by transcriptome data. ** Endoreduplication is absent in the vegetative body but reported in the megagametophyte of Ginkgo and the endosperm of rice. Important sequences are highlighted in bold. (b) Unrooted Maximum-likelihood phylogenetic tree of SMR homologues. The complete tree is available at Supplemental File 2. (c) Multiple sequence alignment showing conservation of the three characteristic SMR motifs (A, B and C) in Chara braunii (Cb), Klebsormidium nitens (Knit), Nitella mirabilis (Nm), *Marchantia polymorpha (Mp), Arabidopsis thaliana (AT),* and *Physcomitrium patens (Pp).* Amino acids are coloured according to the Taylor scheme. Consensus sequence is shown at the bottom.

Phylogenetic analysis recovered well-supported clades that largely mirrored the accepted relationships among major land-plant lineages (Figure 2B, Supplemental File 2), supporting the identification of the recovered sequences as bona fide SMR homologues rather than spurious HMM matches. Bryophyte SMRs formed a distinct clade, with a marked expansion in mosses but single-copy genes in most liverworts and hornworts. Additional lineage-specific clades were recovered in lycophytes, ferns and seed plants, indicating that the SMR family expanded independently several times during land-plant evolution. Despite these independent expansions, SMRs were conserved in all land-plant lineages surveyed, and neither copy number nor family expansion correlated with the occurrence of vegetative endoreduplication.

All recovered proteins, including CbSMR, retained the three conserved motifs (A, B and C) that define the SMR family (Kumar et al., 2015)(Figure 2C). Motif A is the CDK-binding motif and mediates interaction with cyclin–CDK complexes (Wang et al., 2020a); Motif B is a proline-rich basic region; and Motif C is a putative cyclin-binding motif. All three were identifiable in the *C. braunii* sequence, despite it being the most divergent protein in our phylogeny. The presence of the complete motif architecture in CbSMR suggests that the characteristic SMR motif architecture was already established in streptophyte algae, before the origin of land plants.

### Mp*SMR* is dynamically expressed during development

The genome of *M. polymorpha* contains a single SMR gene (Mp*SMR*, *Mp1g14080*), providing a simplified system to examine the role of SMRs in a non-seed plant that lacks endoreduplication. To examine the developmental regulation of MpSMR, we analysed publicly available transcriptomic datasets covering major developmental transitions. Mp*SMR* transcripts were detected throughout development but were dynamically regulated: expression increased during sporeling germination and was transiently induced during the early stages of regeneration.

To examine spatial regulation, we generated a *_pro_*_M_p*SMR* transcriptional reporter. Reporter activity was detected primarily in peripheral regions of the growing thallus and in mature differentiated cells, distinct from the EdU-positive domains observed in wild-type plants (Figure 3c), consistent with a role in limiting cell proliferation. This expression pattern resembles that of SIM, SMR1, SMR4, and SMR11 in Arabidopsis (Yi et al., 2014).

**Figure 3.**
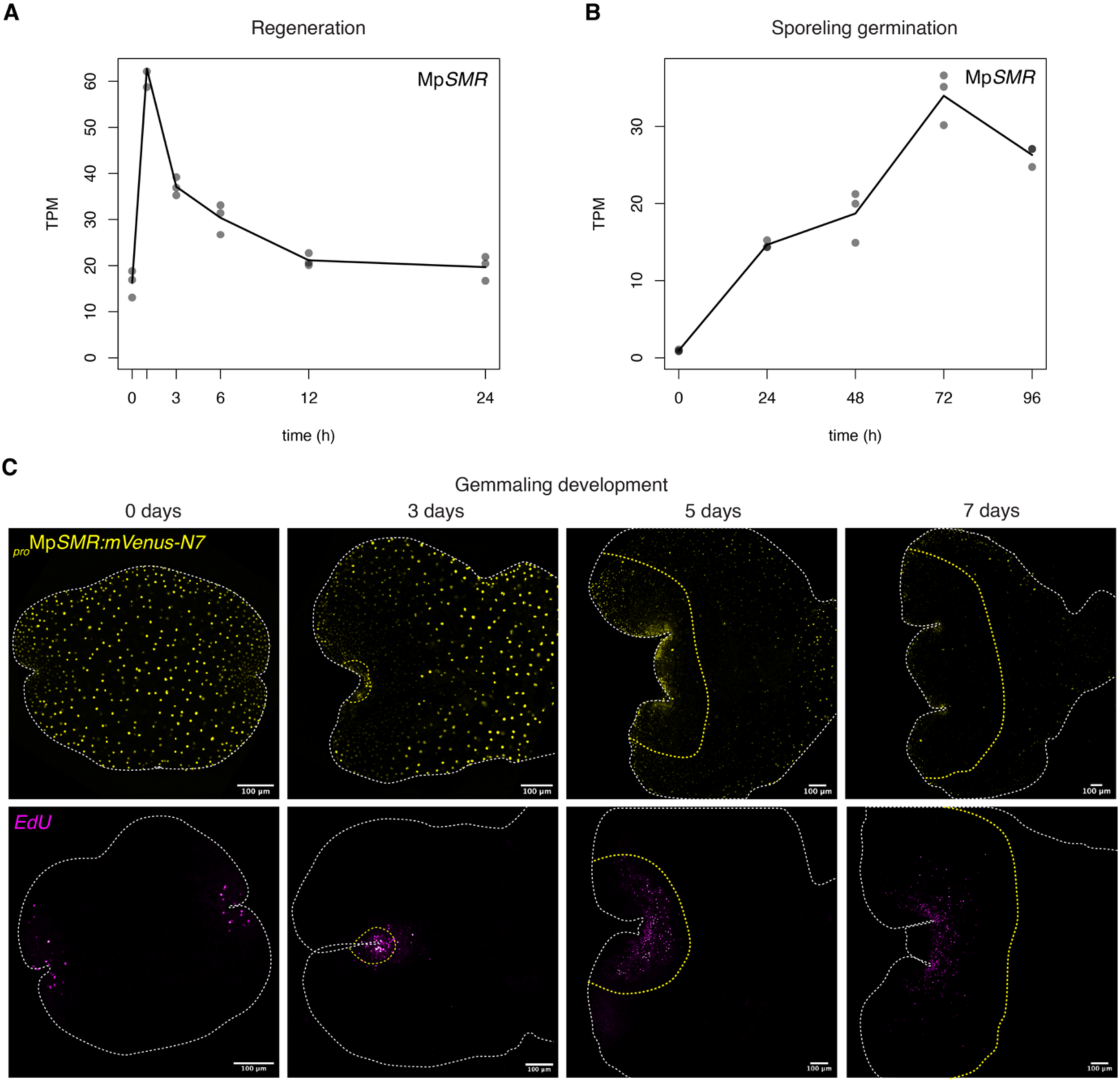
Spatiotemporal expression of MpSMR in Marchantia polymorpha. (a-b) RNA-seq quantification of MpSMR expression (transcripts per million, TPM) during thallus regeneration (a) and sporeling germination (b). Regeneration time course: 0, 1, 3, 12 and 24 h after fragmentation (n = 3 biological replicates; error bars = ± SD). Sporeling germination time course: 0, 24, 48, 72 and 96 h post-plating (n = 3; ±SD). (c) Confocal projections of gemmalings expressing the transcriptional reporter _proM_pSMR:Venus-N7. EdU (5-ethynyl-2’-deoxyuridine) incorporation (magenta) of a wild-type gemmalings at corresponding time points is shown to indicate regions of cell proliferation. Cell outlines (dashed white lines) delineate thallus margins and meristematic domains; yellow dashed lines indicate the apical notch. Scale bars, 100 µm.

Together, these observations show that Mp*SMR* is developmentally and spatially regulated, with transient induction during regeneration and stable expression in non-dividing cells, consistent with a role in limiting cell proliferation.

### MpSMR functions as a conserved cell-cycle inhibitor

We previously showed that the ectopic expression of Mp*KRP* causes enlarged cell size and reduced growth rate (Romani et al., 2026). To determine the function of MpSMR in Marchantia we first generated heat-shock–inducible overexpression lines for MpSMR together with equivalent lines for the single MpKRP. Following induction, both MpKRP and MpSMR strongly inhibited cell proliferation and promoted cell expansion, producing smaller thalli with cells up to 10-fold larger (Figure 4a–b), consistent with the effects of SMR overexpression previously reported in Arabidopsis (Churchman et al., 2006). Cell differentiation was unaffected, and normal growth resumed once induction ceased. The MpSMR fusion protein localised to the nucleus, consistent with a role in modulating CDK activity (Figure 4d) (Churchman et al., 2006).

**Figure 4.**
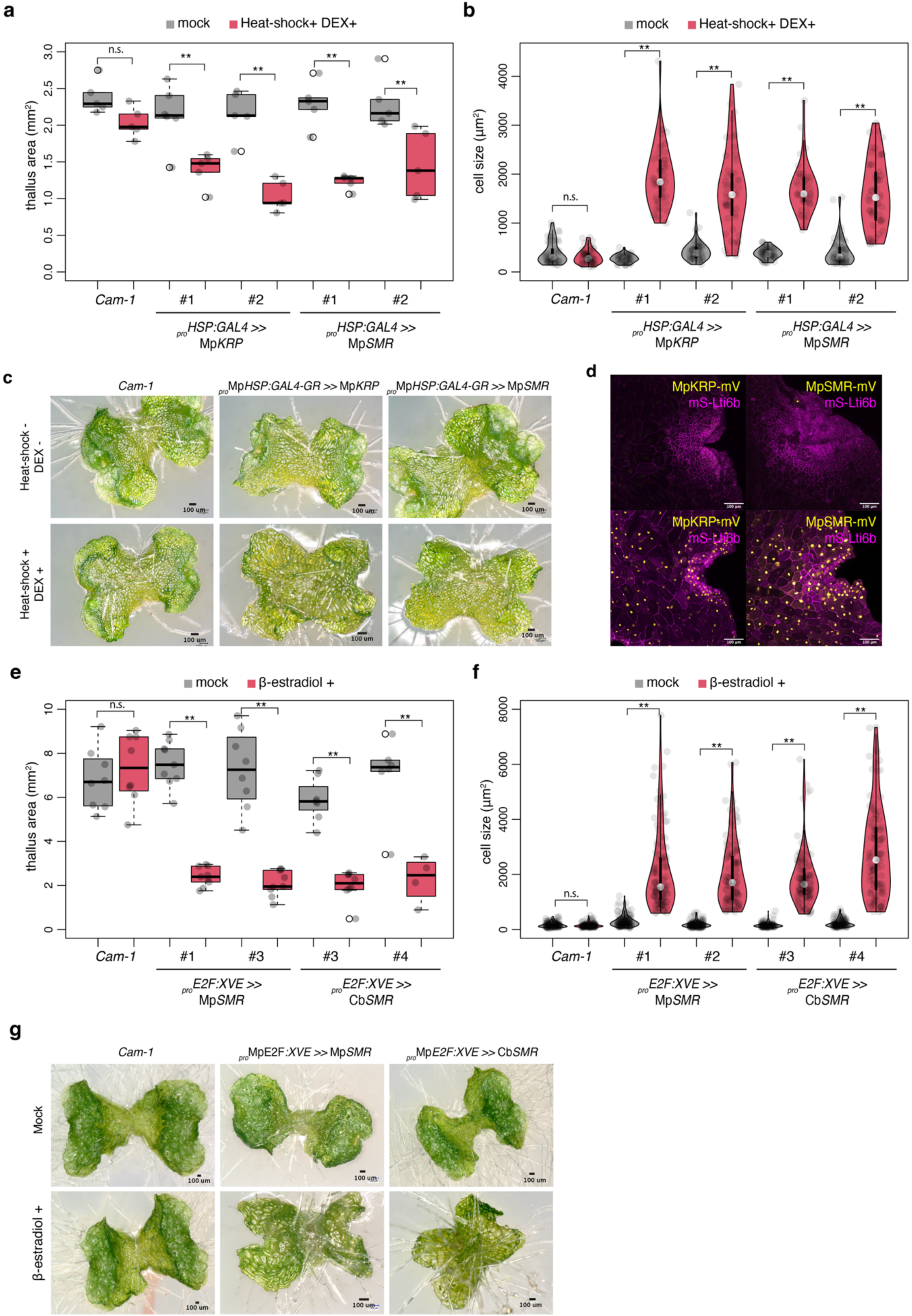
*Functional consequences of MpKRP, MpSMR and CbSMR overexpression in* Marchantia polymorpha. (a–b) Quantification of thallus area (a) and cell size (b) of 4-day-old gemmalings following heat-shock induction or mock treatment of *_proH_SP17.8:*Mp*KRP–mVenus* (#1 and #2 independent lines) and *_proH_SP17.8:*Mp*SMR–mVenus* (#1 and #2 independent lines). (c) Representative bright-field images of plants corresponding to (a–b). (d) Representative confocal maximum-intensity projections of *_proH_SP17.8:* Mp*KRP–mVenus* and *_proH_SP17.8:* Mp*SMR–mVenus* lines following heat-shock induction or mock treatment. Nuclear-localised mVenus is shown in yellow. Plasma membrane signal corresponds to a constitutive *_pro_*_M_p*UBE2:mScarlet-Lti6b* in magenta. (e–f) Quantification of thallus area (e) and cell size (f) of 7-day-old gemmalings following β-estradiol induction or mock treatment of *_pro_*_M_p*E2F:XVE >>* Mp*SMR–mVenus* (#1 and #3 independent lines) and *_pro_*_M_p*E2F:XVE >>* Cb*SMR–mVenus* (#3 and #4 independent lines). (g) Representative bright-field images of plants corresponding to (e–f). Scale bars: 100 µm.

**Figure 5.**
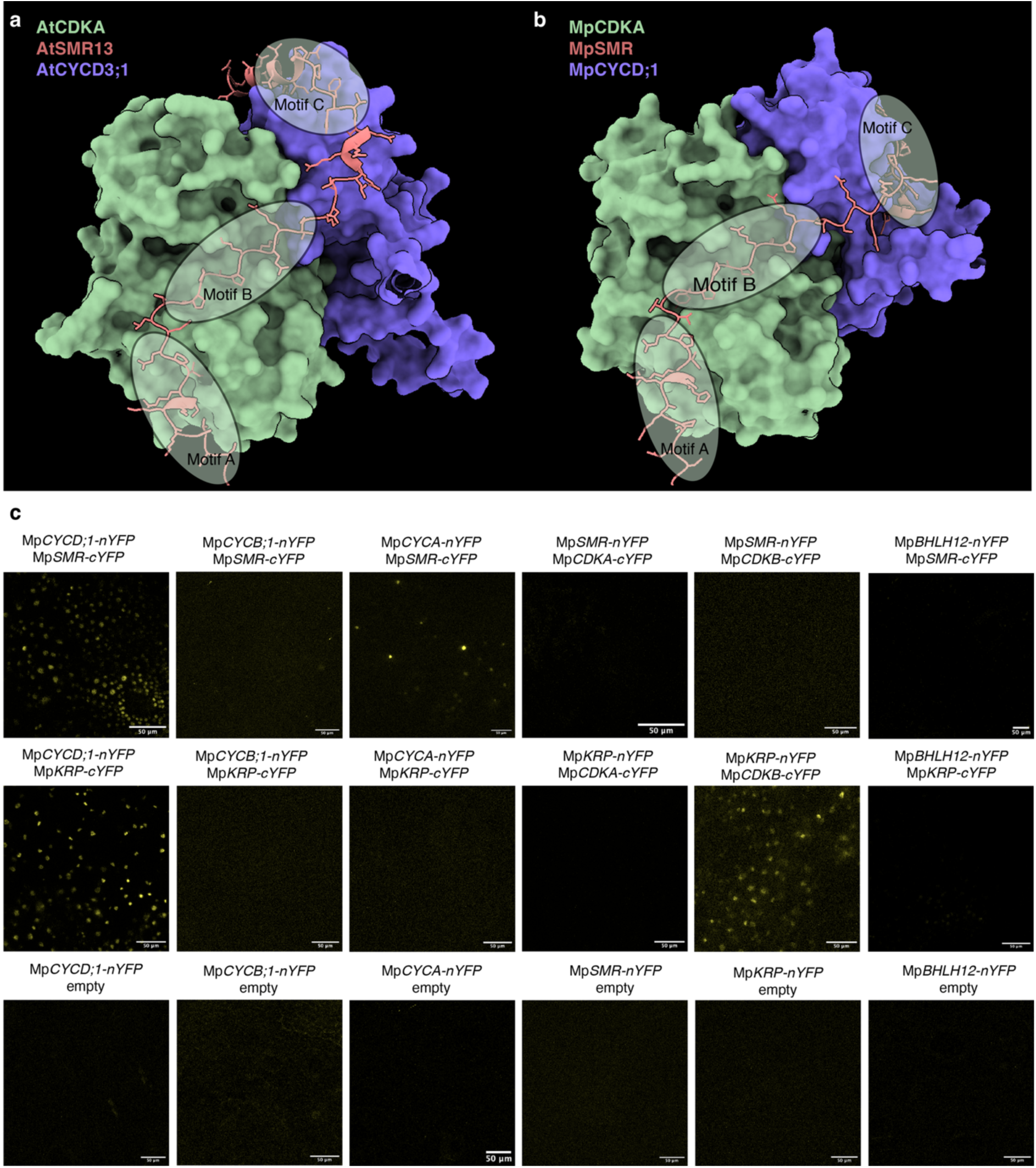
Protein-protein interactions of Marchantia CKI. (a–b) AlphaFold 3 predictions of the Arabidopsis AtCDKA–AtSMR13–AtCYCD3;1 (a) and Marchantia MpCDKA–MpSMR–MpCYCD;1 (b) complexes. SMR (red), CYCD (purple) and CDKA (green) proteins are depicted in different colours. Conserved SMR motifs A–C are highlighted. (c) Bimolecular fluorescence complementation (BiFC) assays in Marchantia polymorpha sporelings. MpBHLH12 served as a negative control. Scale bars = 50 μm.

We repeated this experiment using the estradiol-inducible (XVE) system (Ishida et al., 2022). Consistent with previous results, MpSMR produced smaller thalli with larger cells as seen under heat shock (Figure 4e-f). DAPI staining followed by flow-cytometric analysis revealed a single 1C peak in both induced and control plants, consistent with previous observations in wild-type plants (Nishihama et al., 2015). This supports the functional similarity between MpSMR and MpKRP (Romani et al., 2026), showing that MpSMR promotes cell enlargement without detectable changes in ploidy.

As we also found homologues in streptophyte algae, we heterologously expressed the high-confidence SMR homologue from the charophyte alga *Chara braunii* (CbSMR) in Marchantia. Interestingly, CbSMR also phenocopied the effect of MpSMR, producing cells of comparable size (Figure 4e–f). Constitutive expression of CbSMR (*_pro3_5S:*Cb*SMR*) also led to plants with enlarged cells (Figure S2). The ability of both MpSMR and CbSMR to suppress cell proliferation and promote cell enlargement in Marchantia suggests that this SMR activity predates the emergence of land plants and endoreduplication.

### MpSMR retains conserved interactions with the G1 cyclin–CDK complex

In angiosperms, SMR and KRP proteins interact with cyclin–CDK complexes to regulate cell-cycle progression. In Marchantia, the G1 cyclin–CDK complex comprises MpCYCD;1 and MpCDKA (Romani et al., 2026). To investigate the interactions of MpSMR with core cell-cycle regulators, we predicted its association with this complex using AlphaFold 3 (Abramson et al., 2024) and tested the interactions of MpSMR and MpKRP in vivo by bimolecular fluorescence complementation (BiFC) in Marchantia sporelings.

AlphaFold 3 predicted MpSMR bound to the assembled MpCDKA–MpCYCD;1 complex with medium confidence (ipTM = 0.65). To assess the conservation of this interaction architecture, we compared the predicted Marchantia complex with the corresponding Arabidopsis complex. We chose AtSMR13 because it showed the highest sequence similarity to MpSMR. Both models showed an analogous spatial arrangement of the conserved SMR motifs A–C: motif A contacted the CDK subunit, motif C contacted the cyclin, and motif B was positioned at the interface between the two subunits. Notably, the positioning of motif B was particularly similar between the two structures, suggesting conservation of the overall SMR interaction architecture.

In BiFC assays, both MpSMR and MpKRP interacted with MpCYCD;1, but not with MpCDKA when tested individually. However, their interaction profiles differed for other cell-cycle regulators. MpSMR also interacted with MpCYCA, whereas MpKRP interacted with MpCDKB. Neither protein showed detectable interaction with MpCYCB1;1. Neither MpBHLH12, used as a negative control, nor the empty-vector controls produced detectable signal.

Together, these results support a conserved interaction of SMR and KRP proteins with the G1 cyclin–CDK machinery, while revealing differences in their interactions with other cyclins and CDKs. These distinct interaction profiles suggest that MpSMR and MpKRP may have overlapping but distinct roles in regulating cell-cycle progression.

### MpSMR loss-of-function reveals a conserved role in growth modulation

To assess the endogenous role of SMRs during vegetative development, we generated CRISPR–Cas9 knockout lines targeting the single MpSMR locus, obtaining two independent alleles carrying insertions that introduced frameshifts and premature stop codons (Figure 6a). Compared with wild type, Mp*smr* mutants showed a slight but significant increase in thallus area and a reduction in average cell size (Figure 6b-d). Because the thalli were larger yet composed of smaller cells, this phenotype is consistent with increased cell proliferation, opposite to the reduced division and cell enlargement observed in the overexpression lines. Together, the reciprocal gain- and loss-of-function phenotypes establish MpSMR as a negative regulator of cell proliferation that contributes to cell-size control.

**Figure 6.**
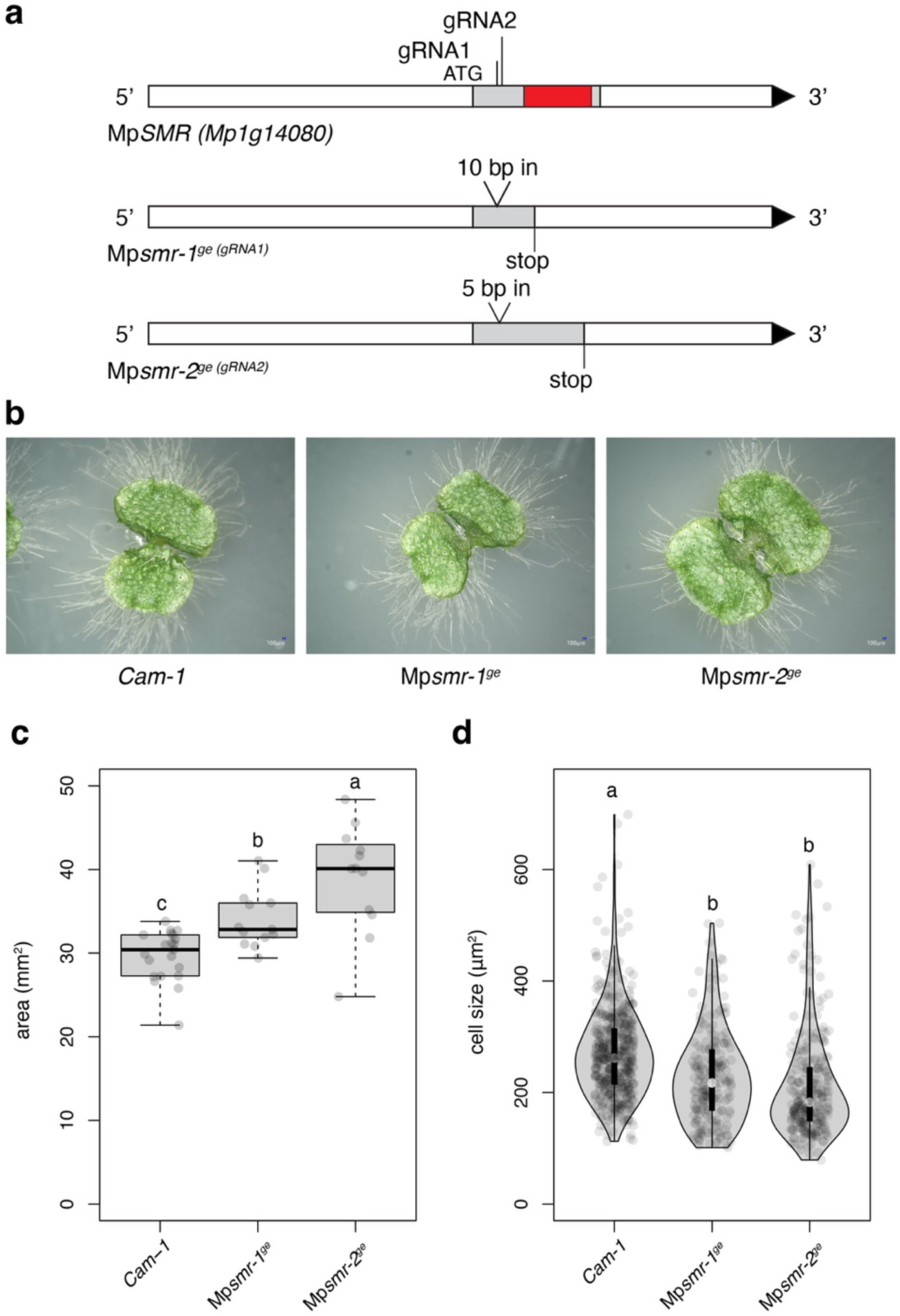
Phenotypes of MpSMR loss-of-function mutants in Marchantia polymorpha. (a) Schematic representation of the MpSMR genomic locus and two CRISPR/Cas9-induced alleles, Mpsmr-1 and Mpsmr-2. Coding sequences (grey boxes), conserved CKI domain (red box), gRNA target sites and mutations are indicated. The 10-bp and 5-bp insertions and resulting premature stop codons are shown. (b) Representative bright-field images of 10-day-old wild-type (Cam-1) and Mpsmr mutant thalli. Scale bars, 100 µm. (c) Quantification of whole-thallus projected area for wild-type and Mpsmr mutants at 10 days post-gemma plating (n = 12 thalli per genotype; boxes: 25th–75th percentiles; whiskers: 1.5× IQR; circles: outliers). (d) Violin plots of individual epidermal cell size (µm²) measured from apical notch regions of wild-type and Mpsmr plants (n ≥ 250 cells per genotype; white dots: median; black bars: interquartile range). Statistical significance (ANOVA followed by Tukey’s HSD test) is indicated by different letters (P < 0.05).

Despite these changes in growth and cell size, Mpsmr mutants showed no major morphological abnormalities: rhizoid formation, air-pore architecture and oil-body patterning were indistinguishable from wild type, suggesting that MpSMR is largely dispensable for vegetative cell differentiation. This subtle but reproducible phenotype parallels the modest growth effects of CKI loss-of-function in other systems, including higher-order SMR mutants in Arabidopsis (Yamada et al., 2022) and the single MpKRP knockout in Marchantia (Romani et al., 2026).

## DISCUSSION

SIAMESE-RELATED (SMR) proteins are a plant-specific family of cyclin-dependent kinase inhibitors that have been primarily studied in Arabidopsis thaliana, where they promote endoreduplication and cell enlargement (Churchman et al., 2006). This functional framework has often been generalized to land plants as a whole. Here we revised this assumption, showing that vegetative endoreduplication is not a universal feature of land plants. The most likely scenario is that endoreduplication was absent from the common ancestor of embryophytes and from that of tracheophytes. In contrast, we found that SMRs predate the origin of land plants and are present in several streptophyte algal lineages, including *C. braunii*. We provide the first *in vivo* functional characterization of an SMR in a non-seed plant, where it controls cell size independently of endoreduplication.

The view that endoreduplication is a core feature of plant growth rests largely on research in flowering plants. Our reconstruction instead indicates that vegetative endoreduplication was absent from the common ancestors of land plants and tracheophytes. Thus, endoreduplication appears to have evolved independently within mosses and along the seed-plant lineage, as previously proposed by Bainard *et a*l. (2020). Given its independent occurrence in animals, the distribution of endoreduplication is best explained by repeated lineage-specific gains rather than an ancient origin followed by widespread loss.

It is important to stress that mosses and angiosperms differ in their dominant generation, the gametophyte and the sporophyte, respectively. In mosses, both generations (the sporophyte and the gametophyte) undergo endoreduplication (Palova et al., 2020). In angiosperms, in addition to the sporophyte, there is evidence of endoreduplication in the female gametophyte and endosperm (Sabelli et al., 2013). Interestingly, endoreduplication also occurs in the megagametophyte of the gymnosperm *Ginkgo biloba* (Avanzi and Cionini, 1971), though not in its sporophyte (Barow and Meister, 2003), raising the possibility that an endocycle in reproductive tissues arose earlier, within spermatophytes.

An important caveat of comparative endoreduplication surveys is that the endocycle could be restricted to rare, highly specialized terminal cell types such as trichomes (De Veylder et al., 2011). An alternative explanation for the observed distribution is that endoreduplication is widespread in some lineages but restricted to specific cell types in others. Similar to angiosperms, in mosses, endoreduplication is observed even in the protonema stage, where cell populations are homogeneous with limited differentiation (Palova et al., 2020), suggesting that endoreduplication occurs with high frequency and not necessarily in differentiated cells. Although detecting endoreduplication in very small cell populations by flow cytometry can be challenging, cell-type-specific endoreduplication has not been conclusively demonstrated in bryophytes. In *Sphagnum*, large hyaline cells have been proposed to undergo endoreduplication (Bainard et al., 2020). However, because these cells later undergo programmed cell death, any such endoreduplication may be transient (Liu et al., 2024). In Marchantia, specialised terminal cell types, such as oil-body cells and rhizoids, could also be candidates for cell-type-specific endoreduplication. In particular, the elongated rhizoid cells may undergo this process. Interestingly, we did not detect defects in the differentiation traits examined in Marchantia Mp*smr* or Mp*krp* mutants (this study; Romani et al., 2026), consistent with endoreduplication not being required for these traits. Thus, the recruitment of SMRs into cell-differentiation programmes may represent a derived function that evolved independently in different plant lineages.

If widespread endoreduplication in plants is a derived feature and might not be associated with specific cells, what were SMR proteins doing before it evolved? Notably, SMR genes are retained across all sampled land-plant lineages, indicating that the absence of vegetative endoreduplication in these lineages cannot be explained simply by the loss of SMR genes. The conserved role of MpSMR in cell size control mirrors what we described for MpKRP (Romani et al., 2026). We also found that both CKIs interact with MpCYCD;1 in planta, consistent with their association with the G1 cyclin–CDK machinery. MpSMR and MpKRP also showed distinct interaction profiles with other cyclins and CDKs. Comparable interactions were reported for PpSMR12 in *Physcomitrium patens* (Kumar et al., 2015). PpSMR12 also complements the Arabidopsis *siamese* mutant when heterologously expressed. The SMR of *C. braunii* arrested division and enlarged cells when expressed in Marchantia, indicating substantial conservation at the protein level and showing that the ancestral SMR was already a functional CKI in a streptophyte alga, before the emergence of land plants.

These findings suggest that an ancestral function of plant CKIs, paralleling that of animal Cip/Kip proteins, was to repress cell-cycle progression independently of endoreduplication. The role of CKIs in endoreduplication cannot be separated from the diversification of other components of the machinery that allows it. In Arabidopsis, SIM promotes endoreduplication by inhibiting mitotic CDK activity, via the G2/M-specific CDKB1;1-CYCA2;3 complex (Boudolf et al., 2009; Wang et al., 2020a). Such complexes may not have existed in the common ancestor of land plants, which may have had a simpler cyclin complement, as observed in Marchantia (Romani et al., 2026). The later diversification of these components during the evolution of seed plants may have enabled the recruitment of CKIs into endocycle control independently in different plant lineages. In that sense, the functional evolution of SMR genes mirrors the independent recruitment of Cip/Kip genes into endocycle control in animals (Ullah et al., 2009).

Regarding the biological role of MpSMR in Marchantia, mutant phenotypes suggest that MpSMR contributes to cell-size regulation. This function may also be relevant to environmental responses, particularly DNA damage. It was recently shown that Mp*SMR* expression is induced upon DNA damage (Li et al., 2026). This is consistent with its role arresting the cell cycle, as this facilitates DNA repair to maintain genome integrity (Yi et al., 2014; Trujillo-Esquivel et al., 2026). This potential ancestral function of SMR proteins in DNA-damage could work independently of endoreduplication as discussed above. However, further research is needed to understand how this response operates in bryophytes. Recent studies indicate that some components of the DNA-damage response are conserved across land plants, whereas others may have diverged (Li et al., 2026). More broadly, our study underscores how conserved cell-cycle regulators can acquire new developmental functions during land-plant evolution and in different cellular contexts.

## ACKNOWLEDGEMENTS

This work was funded by BBSRC BB/T007117/1 to J.H., and BBSRC BB/F011458/1 for confocal microscopy. F.R. is a Leverhulme Early Career Fellow (ECF-2023-534) funded by the Leverhulme Trust and the Isaac Newton Trust (23.08(f)). We thank Miguel A Blazquez and Antonio Serrano Mislata (IBMCP, Valencia, Spain) for sending CRISPR constructs to target Mp*SMR* and for helpful discussion.

## MATERIAL AND METHODS

### Ancestral-state reconstruction

Flow-cytometry datasets from published literature (Barow and Meister, 2003; Bainard and Newmaster, 2010; Bainard et al., 2011; Bainard et al., 2012; Bainard and Villarreal, 2013; Bainard et al., 2020) were compiled to score EI values and the presence or absence of endoreduplication across land plant lineages as shown in Supplemental Table S1. Only vegetative tissues were included. When more than one value was available, the average was assigned to the taxa.

Ancestral states were reconstructed on a phylogeny of the 202 sampled taxa using available trees as references. Endoreduplication index was treated as a continuous character, and ancestral values were estimated by squared-change parsimony (Maddison, 1991). Alternatively, endoreduplication was treated as a binary presence/absence character (EI ≥ 0.1), and ancestral states were reconstructed in Mesquite v4 (https://www.mesquiteproject.org) using either marginal-probability reconstruction under a one-parameter Markov k-state model (Lewis, 2001) or maximum-likelihood reconstruction. Nodes with a reconstructed EI < 0.1 were interpreted as non-endoreduplicating.

Additionally, ancestral states were reconstructed for the same tree using the ancestral.pars function implemented in the R package phangorn (Schliep, 2011). Endoreduplication index (EI) was treated as a continuous character, and ancestral states were estimated using squared-change parsimony with the POSTORDER method.

### Phylogenetic identification and analysis of SMR proteins

Proteomes from representative eukaryotic lineages were obtained from Phytozome, Phycocosm, EnsemblPlants, and species-specific genome portals (Goodstein et al., 2012; Bolser et al., 2017; Grigoriev et al., 2021). The dataset included the following species: *Chlamydomonas reinhardtii (Merchant et al., 2007), Ostreococcus lucimarinus (Palenik et al., 2007), Klebsormidium nitens (Hori et al., 2014), Mesostigma viride (Liang et al., 2020), Chlorokybus atmophyticus (Wang et al., 2020b), Chara braunii (Nishiyama et al., 2018), Penium margaritaceum (Jiao et al., 2020), Zygnema cylindricum (Feng et al., 2024), Mesotaenium endlicherianum (Cheng et al., 2019), Spirogloea muscicola (Cheng et al., 2019), Spirogyra pratensis (Goldbecker et al., 2026), Anthoceros agrestis Oxford* (Li et al., 2020a)*, Phaeoceros laevis (Li et al., 2020b), Marchantia polymorpha* v6.1 (Montgomery et al., 2020)*, Lunularia cruciata (Linde et al., 2021), Physcomitrium patens* v6.1 (Bi et al., 2024), *Ceratodon purpureus (Carey et al., 2021), Sphagnum fallax (Healey et al., 2023)(X), Takakia lepidozioides* (Hu et al., 2023)*, Diphasiastrum complanatum (Xu et al., 2024), Huperzia asiatica (Li et al., 2024), Selaginella moellendorffii (Banks et al., 2011), Salvinia cucullata (Li et al., 2018), Ceratopteris richardii v2.1* (Marchant et al., 2022)*, Pteridium aquilinum (Suissa et al., 2026), Thuja plicata (Shalev et al., 2022), Ginkgo biloba v2* (Liu et al., 2021)*, Amborella trichopoda* v1.0 (Amborella Genome, 2013)*, Setaria viridis (Mamidi et al., 2020), Oryza sativa (International Rice Genome Sequencing, 2005), Arabidopsis thaliana (Arabidopsis Genome, 2000), Solanum lycopersicum (Tomato Genome, 2012).* Additionally, the transcriptomes of charophyte algae were obtained from Cooper and Delwiche (https://figshare.com/articles/dataset/Green_algal_transcriptomes_for_phylogenetics_and_comparative_genomics/1604778).

Putative SMR homologues were identified using the InterPro HMM profile IPR040389 and hmmsearch from HMMER v3.4 (Finn et al., 2011). Searches were performed on the Galaxy server (usegalaxy.eu) (Galaxy, 2026). Only high-confidence hits (E-value < 1e–5) were retained for subsequent analyses. Sequences were retrieved using the SeqinR package in R (Charif and Lobry, 2007).

Protein sequences were aligned using MAFFT in Jalview (Procter et al., 2021) with default options except for gap open penalty = 1 and gap extension penalty = 0.05. Alignments were trimmed to retain positions with ≥50% site occupancy for phylogenetic analysis. Maximum-likelihood phylogenies were inferred with IQ-TREE 3 (Minh et al., 2020) with 1000 bootstrap replicates and default parameters. Trees were visualised and rooted using iTOL v6 (Letunic and Bork, 2024). Full phylogenies are provided in Supplemental File 2.

### Expression analysis of MpSMR

Transcript abundance (TPM) across Marchantia polymorpha development—including sporeling germination, vegetative stages, and regeneration—was extracted from MarpolBase. Regeneration RNA-seq datasets (DRR330148–DRR330173) were downloaded from NCBI SRA, mapped to *M. polymorpha* Tak-1 genome v5.1 using HISAT2 with default parameters, and quantified with HTSeq with default parameters. Counts were then normalised to TPM.

### Plasmid construction

The Mp*SMR* coding and promoter sequence were obtained from the *M. polymorpha* Tak-1 genome version 5.1 (Montgomery et al., 2020) and the Cb*SMR* CDS from its reference genome (Nishiyama et al., 2018). The CDSs were manually domesticated by introducing synonymous mutations to remove internal BsaI and SapI sites. The sequences of synthetic L0 parts used in this work are available in Table S1. L0 parts were synthesized by GENEWIZ following the standard syntax for plant synthetic biology, with CDS and PROM5 or PROM and 5UTR overhangs, and cloned into the pUAP1 plasmid (Addgene #63674) (Patron et al., 2015) by homology recombination. Other parts were derived from previous works (Sauret-Gueto et al., 2020; Romani et al., 2026).

The binary acceptor pBy01 (Romani et al., 2024) was used for assembly with L0 parts corresponding to PROM5 or PROM and 5UTR. The pBy12 and pBy20 vectors (Tse et al., 2024) were used for bimolecular fluorescence complementation and constitutive overexpression, respectively, while pBy13 and pBy23 (Romani et al., 2026) were used for the estradiol-inducible system. Plasmids were assembled as detailed in Supplemental Table S1 using Type-IIS cloning as described previously for L3 plasmids (Romani et al., 2024). The presence of the correct insert was confirmed by restriction XhoI digestion (Thermo Scientific #FD0694) and Sanger sequencing.

For heat-shock-inducible expression, the corresponding parts (Supplemental Table S1) were first assembled in a pCk3 vector and subsequently cloned into a pCsA vector using Loop Assembly (Pollak et al., 2019) with parts described in Supplemental Table S1.

For genome editing, CRISPR–Cas9 constructs targeting MpSMR (Mp1g14080) were generated using pMpGE_En03 and then subcloned into pMpGE010 vector as previously described (Sugano et al., 2018). gRNA1 was generated using primers sgRNAMpSMR_1Fw (CTCGGTCGGAGGACATGAGTCAAC), sgRNAMpSMR_1Rev (AAACGTTGACTCATGTCCTCCGAC); and gRNA2 with primers sgRNAMpSMR_2Fw (CTCGATGAGTCAACAGGACACTTG) and sgRNAMpSMR_2Rev (AAACCAAGTGTCCTGTTGACTCAT).

### Plant transformation and growth

Marchantia polymorpha *Cam-1* (male) and *Cam-2* (female) accessions were used unless otherwise stated. Plants were grown in FitoClima 1200PL cabinets (Aralab) on half-strength Gamborg B5 basal medium (Phytotech #G398) at pH 5.8 with 1.2% agar micropropagation grade (Phytotech #A296) and 0.5 % (w/v) sucrose (Sigma-Aldrich #16104) under continuous LED light (90 µmol m⁻² s⁻¹) at 21°C. For spore production, plants were grown in Microbox micropropagation containers (SacO2) in long-day conditions (16 h light/8 h dark) under light supplemented with far-red light as described (Sauret-Gueto et al. 2020).

Stable transformation of spores was performed using *Agrobacterium tumefaciens* GV3101 via a modified spore transformation method (Annese et al., 2025). Transformants were selected on hygromycin (20 µg/mL) or chlorosulfuron (0.5 µM). For CRISPR–Cas9 knockout lines, thallus transformation followed the method described by Kubota et al. (Kubota et al., 2013).

At least four independent lines were screened for each fluorescent reporter, and the consensus expression pattern is shown.

### Phenotyping and microscopy

Thallus area was quantified from images of gemmalings from 4-10 days old using ImageJ. For cell-size measurements, cells surrounding the dividing and differentiating zone (DDCZ) were manually segmented from 200× images, excluding the small meristematic region. At least 50 cells per plant were measured, with ≥3 biological replicates.

Representative images of Marchantia plants were taken using a Keyence VHX-5000 digital microscope equipped with a 20x-200x Ultra-Small, High-Performance Zoom Lens (VH-Z20R/Z20T). For β-estradiol (Merck #E8875) treatments, a 50 mM stock solution was dissolved in DMSO and added to the molten Gamborg B-5 agar medium at a final concentration of 5 μM as described in Ishida et al. (Ishida et al., 2022). Plants carrying the heat-shock promoter were grown under normal conditions or on medium supplemented with 5 μM dexamethasone (#265005, Sigma-Aldrich) and incubated in a 37 °C oven for 2 h daily before being returned to standard growth conditions. Plants were imaged after 4 days.

Confocal imaging used a Leica SP8X system with white-light and 405/442 nm lasers. mVenus excitation/emission was set to 514 nm / 527–552 nm. Chlorophyll autofluorescence was captured at 633 nm / 687–739 nm. Sequential scanning was used for fluorophores with overlapping spectra.

### EdU labeling

Gemmalings were grown in normal conditions for 1, 3, 5, or 7 days and incubated in liquid half-strength Gamborg B-5 medium under continuous light for 3 h with 20 μM 5-ethynyl-2′-deoxyuridine (EdU) from the Click-iT EdU Imaging kit with Alexa Fluor 488 (Invitrogen #C10337). Then they were fixed with 4% formaldehyde for 1 h and washed twice with phosphate buffer saline (PBS) and 0.5% Triton X-100 in PBS for 20 min. After washes, samples were incubated with a freshly prepared reaction mixture following the manufacturer’s instructions but without the Hoechst 33,342 component. After labeling, samples were protected from light, washed twice with PBS and soaked in iTomei-D (Tokyo Chemical Industry #T3940) clearing solution and mounted in 70% w/v iohexol (Tokyo Chemical Industry #I0903) in PBS as previously described (Sakamoto et al. 2022). Samples were covered with a glass coverslip and imaged on the Leica SP8X confocal microscope as described before (Romani et al., 2026).

### Flow cytometry

Nuclear DNA content was measured in thallus tissue of proMpE2F:XVE >> MpSMR–mVenus and wild-type (Cam-1) plants grown for 7 d and then treated with 5 µM β-estradiol or mock solution for a further 10 d. Nuclei were released by chopping 50–100 mg of fresh thallus with a razor blade in ice-cold Galbraith’s buffer (Galbraith et al., 1983) and stained with DAPI as described by Nishihama et al. (2015).

Nuclei were analysed using a CytoFLEX flow cytometer (Beckman Coulter). DAPI was excited using the 405 nm laser and collected through a 450/45 nm bandpass filter; 660/10 nm emission from the 638 nm laser was recorded in parallel and used with the DAPI signal to gate intact nuclei and exclude chloroplast-derived particles and debris. Data was analyzed using Floreada (https://floreada.io/).

### Protein-protein interactions

To predict the structure of the interaction between SMR proteins and their cyclin–CDK complexes, full-length protein sequences of MpSMR, MpCDKA and MpCYCD;1 from the M. polymorpha Tak-1 v5.1 genome were submitted to AlphaFold 3 (Abramson et al., 2024) as a ternary complex (MpSMR–MpCDKA–MpCYCD;1) using the AlphaFold Server (https://alphafoldserver.com) with default settings. The same was done for their Arabidopsis counterparts (AtSMR13–AtCDKA–AtCYCD3;1). Protein complexes were visualized using ChimeraX and the top-ranked model is shown.

BiFC was performed as described in Romani et al. (2026). Coding sequences of MpSMR, MpKRP, MpCYCD;1, MpCYCA, MpCYCB;1, MpCDKA and MpBHLH12 (Kongsted et al., 2026) were fused to the N- or C-terminal half of YFP as indicated, and co-transformed in *M. polymorpha* sporelings by Agrobacterium infiltration using two antibiotics. 14-day-old sporelings that survived selection were imaged to detect reconstituted YFP fluorescence on a Leica SP8X confocal microscope (excitation 514 nm, emission 527–552 nm), using identical acquisition settings across all combinations.

### Statistics

Statistical analyses were performed in R v4.4.1. Differences between experimental groups were assessed using analysis of variance (ANOVA), followed by Tukey’s honest significant difference test (α = 0.05). The experimental unit, sample sizes and number of biological replicates are indicated in the corresponding figure legends.

