## Supplemental Figures for "An ancestral role of SIAMESE-RELATED proteins in cell-division arrest predates the emergence of endoreduplication in land plants"

**Supplemental Figure S1. Flow-cytometric analysis of nuclear DNA content in MpSMR-overexpressing Marchantia plants.** Nuclei were isolated from $_{pro}$MpE2F:XVE >> MpSMR–mVenus plants grown for 7 days and subsequently treated with β-estradiol or mock solution for 10 days. Histograms show nuclear DNA-content distributions, and lower panels show the gating of intact nuclei.

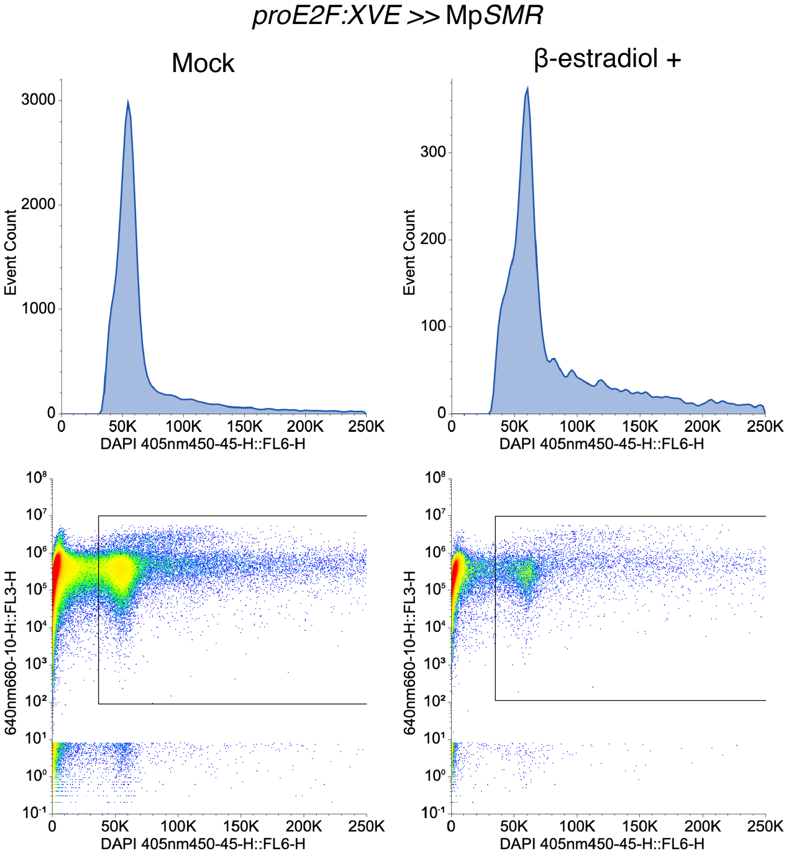


**Supplemental Figure S2. Constitutive expression of CbSMR in Marchantia.** (a) Quantification of epidermal cell size in wild-type Cam-1 and two independent $_{pro}$35S:CbSMR–mVenus lines. (b) Representative bright-field images of wild-type and CbSMR-overexpressing plants at 7 days post-germination. Scale bar is shown in each panel.

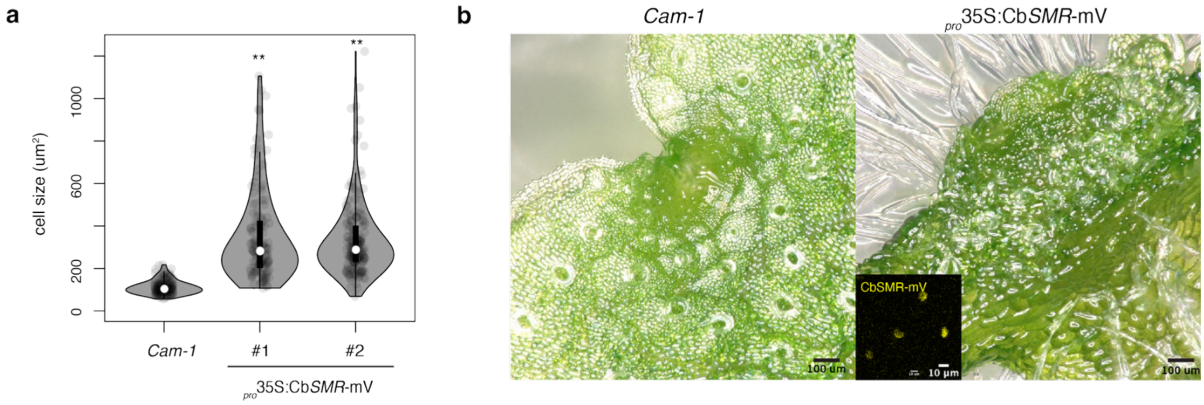


### SUPPLEMENTAL TABLES

**Supplemental Table 1.** Vectors, cloning, primers, and details of gene IDs used.

**Supplemental File 1. Nexus file containing the phylogenetic tree and character data used for ancestral-state reconstruction of endoreduplication in Figure 1.**

**Supplemental File 2.** Complete Unrooted Maximum-likelihood phylogenetic tree of SMR proteins in streptophytes.
