## Supplementary figures and images for "An ancestral role of SIAMESE-RELATED proteins in cell-division arrest predates the emergence of endoreduplication in land plants"

### Supplemental File 2

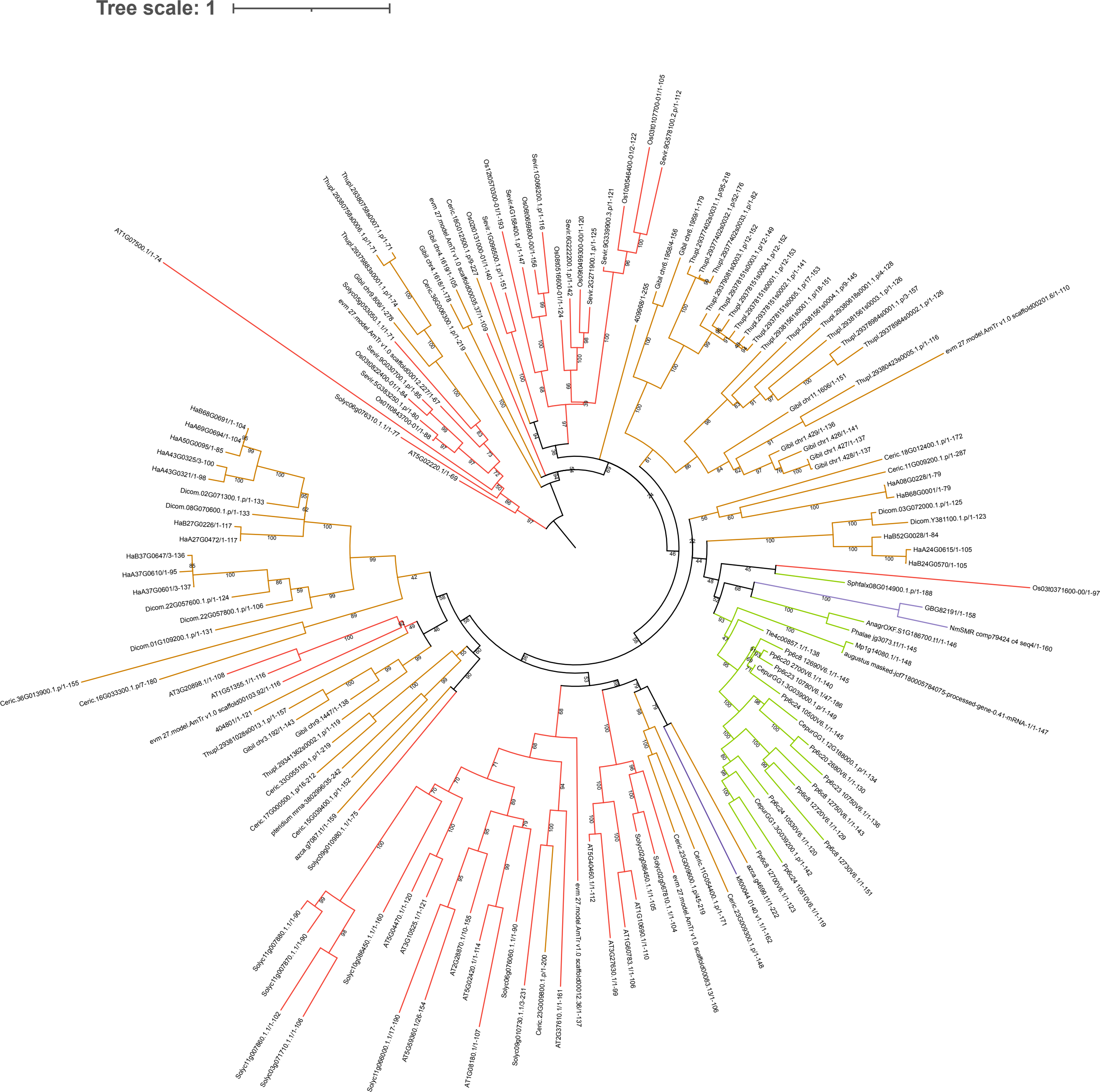
